# Structural basis for recruitment of the CHK1 DNA damage kinase by the CLASPIN scaffold protein

**DOI:** 10.1101/2020.10.02.323733

**Authors:** Matthew Day, Sarah P. Morris, Jack Houghton-Gisby, Antony W. Oliver, Laurence H. Pearl

## Abstract

CHK1 is a protein kinase that functions downstream of activated ATR to phosphorylate multiple targets as part of intra-S and G2/M DNA damage checkpoints. Its role in allowing cells to survive replicative stress has made it an important target for anti-cancer drug discovery. Activation of CHK1 by ATR depends on their mutual interaction with CLASPIN – a natively unstructured protein that interacts with CHK1 through a cluster of phosphorylation sites in its C-terminal half. We have now determined the crystal structure of the kinase domain of CHK1 bound to a high-affinity motif from CLASPIN. Our data show that CLASPIN engages a conserved site on CHK1 adjacent to the substrate-binding cleft, involved in phosphate sensing in other kinases. The CLASPIN motif is not phosphorylated by CHK1, nor does it affect phosphorylation of a CDC25 substrate peptide, suggesting that it functions purely as a scaffold for CHK1 activation by ATR.

## INTRODUCTION

Checkpoint kinase 1, or CHK1, plays a key role in the cellular response to DNA damage at different points in the cell cycle through its function in both the intra S-phase checkpoint, which allows damage to be repaired to facilitate the accurate duplication of genomes, and the G2/M checkpoint, which prevents entry into mitosis of cells with damaged chromosomes (Smits and Gillespie, 2015). Amongst many other targets involved in these two checkpoint responses, CHK1 phosphorylates key residues in the three CDC25 phosphatases, resulting in their deactivation by subsequent degradation or export from the nucleus. This leads to cell cycle arrest as the phosphatases can no longer activate cyclin dependent kinases CDK1 or CDK2, the drivers of the cell cycle. Additionally CHK1 has other important functions in S-phase including a role in origin firing, fork progression, stabilising stalled replication forks, fork restart, and pausing the cell cycle under conditions of replication stress (Iyer and Rhind, 2017).

CHK1 consists of an N-terminal kinase domain (Chen et al., 2000) joined by a flexible linker region to a C-terminal kinase associated (KA1) domain. As an attractive cancer drug discovery target, many crystal structures of the CHK1 kinase domain (CHK1-KD) have been determined bound to various fragments and inhibitors. In all structures of CHK1-KD, whether in apo form or with an inhibitor bound, the kinase domain is in an active conformation with an ordered activation loop, but with a slightly open lobe arrangement leading to the misalignment of the catalytic residues. CHK1 activation occurs when key residues in the linker region are phosphorylated by ATR (Niida et al., 2007). These disrupt intramolecular interactions between the kinase and kinase associated domains (Emptage et al., 2017) and mutations that disrupt the KA1 domain lead to constitutive activity (Gong et al., 2015). Additional autophosphorylation of the KA1 domain further activates the kinase (Gong et al., 2018) and there may be a role for further phosphorylation-driven intermolecular interactions with other factors.

CLASPIN was identified as an interactor of CHK1 in Xenopus extracts (Kumagai and Dunphy, 2000) that was required for the activation of CHK1 in response to DNA replication blocks and stalled replication forks; a function that is conserved in humans (Clarke and Clarke, 2005). The interaction between CHK1 and CLASPIN has also been demonstrated to play a role in the checkpoint response to UV induced DNA damage (Jeong et al., 2003). CLASPIN is required for the ATR dependent phosphorylation of CHK1 (Kumagai et al., 2004) leading to its activation, and furthermore, the ATR-dependent phosphorylation of CHK1 has been demonstrated to be directly stimulated by CLASPIN in an *in vitro* reconstituted system using recombinant human proteins (Lindsey-Boltz et al., 2009).

CLASPIN is a natively unfolded protein (Wright and Dyson, 2015) with three repeated linear motifs that mediate the interaction with CHK1 (Chini and Chen, 2006; Jeong et al., 2003; Kumagai and Dunphy, 2003). These sites in CLASPIN have been reported to be phosphorylated by CHK1 itself (Chini and Chen, 2006) but also through a CHK1-independent route (Bennett et al., 2008) that involves Casein Kinase 1 (CK1) (Meng et al., 2011). More recently an additional pathway has been described in which the three sites are phosphorylated by CDC7 (Yang et al., 2019). Previous studies implicated a site in CHK1-KD, centred around Arg-129, Thr-153 and Arg-162, required for the binding to the three phosphorylation sites in CLASPIN in immunoblot experiments (Jeong et al., 2003).

Here we biochemically characterise the phosphorylation mediated CLASPIN-CHK1 interaction and determine the crystal structure of a complex between CHK1-KD and a phosphorylated CLASPIN peptide. Our results reveal the molecular details of this interaction in the context of CHK1 activation and uncover a new role for an evolutionarily conserved phosphate binding pocket in the kinase domain, which is adapted to multiple functions in different kinase families.

## RESULTS

### A single site in CHK1-KD can bind to the phosphorylated CLASPIN motifs

To characterise the interactions between CHK1 and the three repeated phosphorylation sites in CLASPIN (**Figure 1a**), we synthesised phosphopeptides corresponding to the three different repeats centred on pT916, pS945 and pS982, and measured their binding to CHK1-KD using fluorescence polarisation. All three peptides independently bound the kinase domain with affinities in the high nanomolar range (**Figure 1b**), with the tightest interaction between the kinase domain and the peptide encapsulating the pS945 site. The interaction was abrogated when treated with lambda phosphatase (**Figure 1c**), demonstrating that the interaction is indeed phosphorylation dependent. Mutations in residues in the putative phosphate binding pocket in the kinase domain, also abolished the interaction (**Figure 1d**).

**FIGURE 1.**
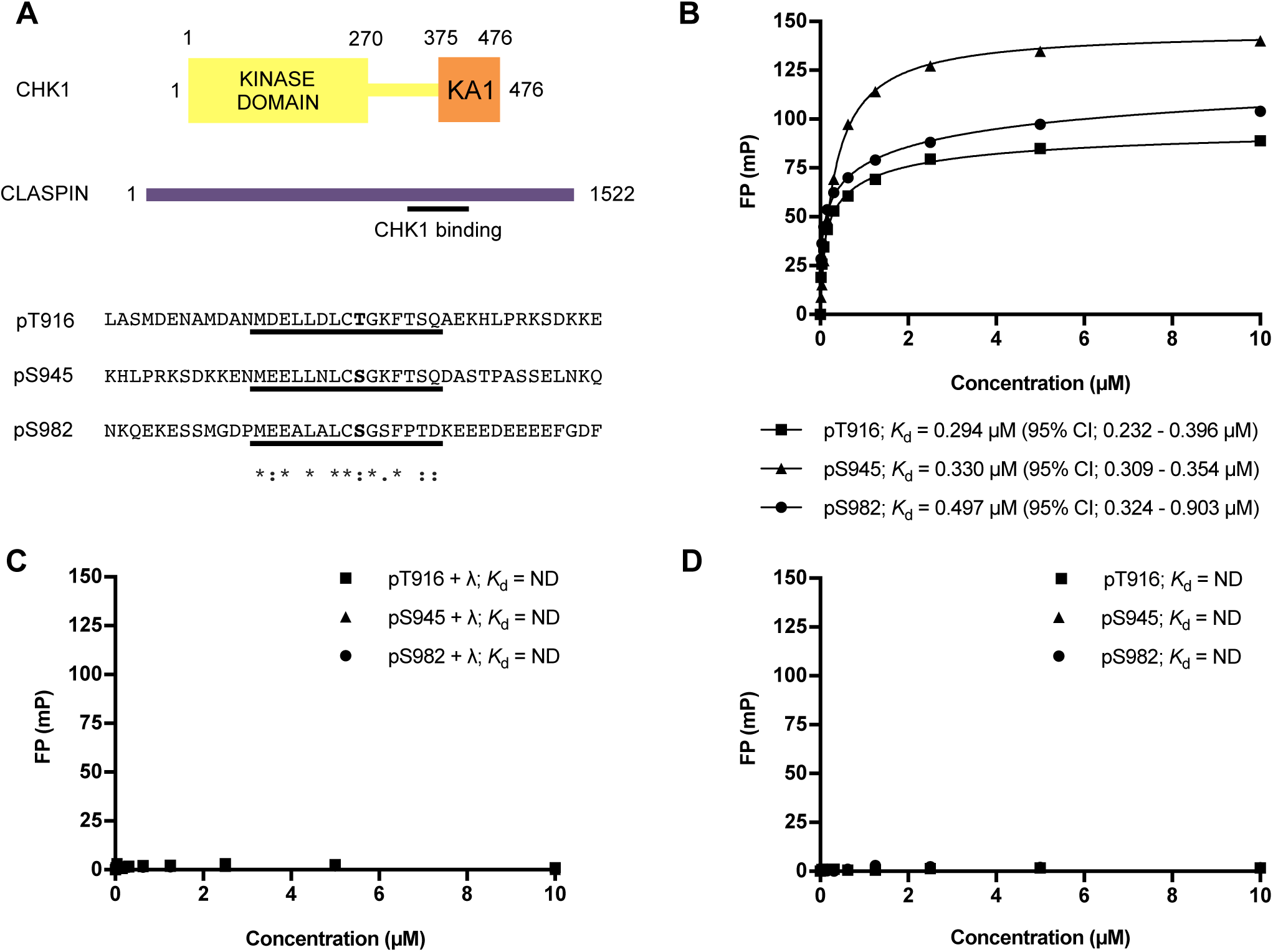
**a)** Schematic representation of CHK1 and CLASPIN and the sequences of the three repeated CHK1 binding motifs in CLASPIN. The underlined regions correspond to the sequences used to synthesize fluorescein labelled peptides and the sequence consensus is shown. **b)** Fluorescence polarisation experiments show binding between a CHK1 construct consisting of only the kinase domain (CHK1-KD) and the three motifs in CLASPIN. **c)** Treatment with λ-phosphatase abolishes binding confirming that the interaction is specific for the phosphorylated CLASPIN peptides. **d)** Mutation of a single site on the kinase domain surface abolishes interaction with the phosphorylated peptides demonstrating that this is the only site responsible for binding to each of the individual motifs in CLASPIN.

### Crystal engineering

Analysis of CHK1-KD structures deposited in the Protein Databank (PDB) reveals two different lattices, in which the protein crystallises (**Figure 2a**). However, attempts to either soak a phosphorylated CLASPIN peptide into crystals of CHK1-KD, or to co-crystallise a complex of CHK1-KD with a CLASPIN peptide, were unsuccessful, yielding crystals of CHK1 alone or in complex with staurosporine in both of the previously observed crystal forms. Examination of the molecular packing (**Figure 2b**) in these crystal forms suggested that the region surrounding the putative phosphate binding site was potentially occluded by contacts with other molecules in the crystal lattice, and this was likely to interfere with binding of a CLASPIN motif in that region (confirmed once the structure of the complex had been determined). In an attempt to relieve this potential restriction, we made a point mutation, distant from both the active site and the putative CLASPIN interaction interface, intended to disrupt a salt-bridge contact in the lattice (**Figure 2c**), thereby destabilizing the predominantly observed crystal form, and allowing new packing variants to be produced.

**FIGURE 2.**
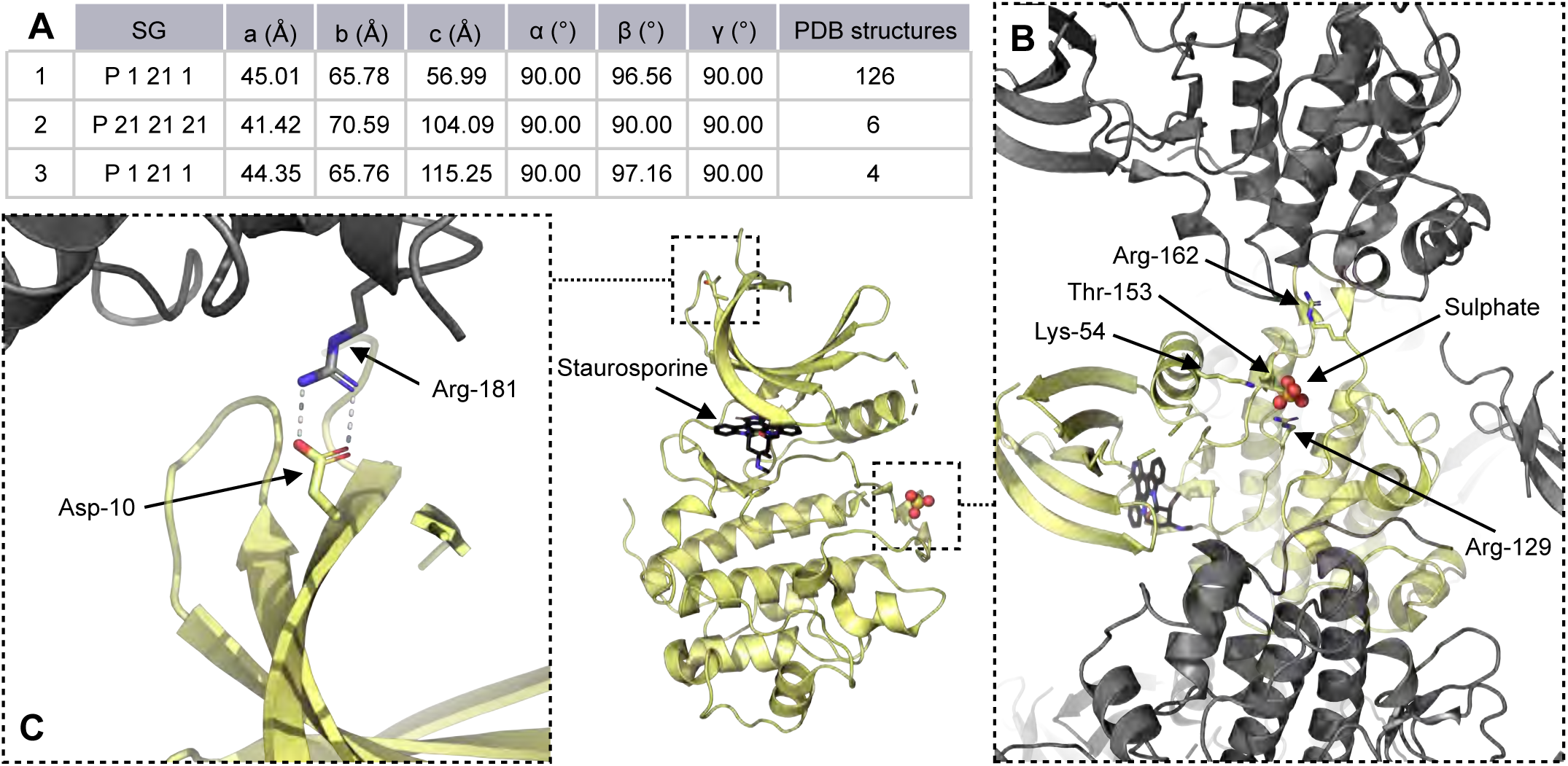
**a)** Summary of CHK1-KD structures in the PDB, cell dimensions are averages across the indicated number of structures in each case. **b)** The asymmetric unit of PDB entry 1NVR, representative of the predominant lattice, contains a single copy of the kinase domain, shown in yellow with symmetry related molecules shown here in grey. The phosphorylated CLASPIN binding site is marked with a sulphate ion in the structure while the active site is occupied by the presence of staurosporine. **c)** Details for the salt bridge between Asp-10 and Arg-181 that is disrupted by the D10R mutation shown alongside its location with respect to the active site and putative CLASPIN interaction surface.

### Crystal structure of CHK1 bound to ATPyS

In an attempt to produce a CLASPIN bound crystal structure of CHK1-KD, various combinations of nucleotides, inhibitors and peptides were combined and set up in crystallisation trials. One of these attempts, where the CLASPIN peptide and ATPyS were included, yielded a structure of the kinase domain bound to the nucleotide alone. The kinase domain is in a similar conformation to that seen in the two known crystal forms, with the nucleotide binding site, activation loop and catalytic residues ordered and positioned in an active, but open, conformation (**Figure 3a**). There is clear density for the ATPγS with the ribose in a C3-endo conformation (**Figure 3b**). This is in contrast to a reported structure of an AMPPNP bound complex in which the ribose was in the C5-endo conformation and the phosphate groups could not be seen (Chen et al., 2000). However, the coordinates and diffraction data for that structure have never been deposited in the PDB, so the reliability of that observation cannot be ascertained. In the ATPγS complex two magnesium ions, chelated by residues Asn-135 and Asp-148, can be seen in the active site balancing the charge on the phosphate groups of the nucleotide. While the phosphate groups can be unambiguously placed in this structure, the γ-phosphate is not positioned in the presumed correct location for catalysis, with the catalytic Asp-130 residue situated ∼8Å from the bridging thioether of the ATPγS (**Figure 3c**). This conformation may be due to constraints imposed by the crystal lattice, or potentially represents a nucleotide bound open conformation formed prior to substrate peptide binding, which might promote full closure of the active site to bring the aspartate residue into the correct position for catalysis to occur. A substrate peptide bound structure would help to distinguish between these possibilities.

**FIGURE 3.**
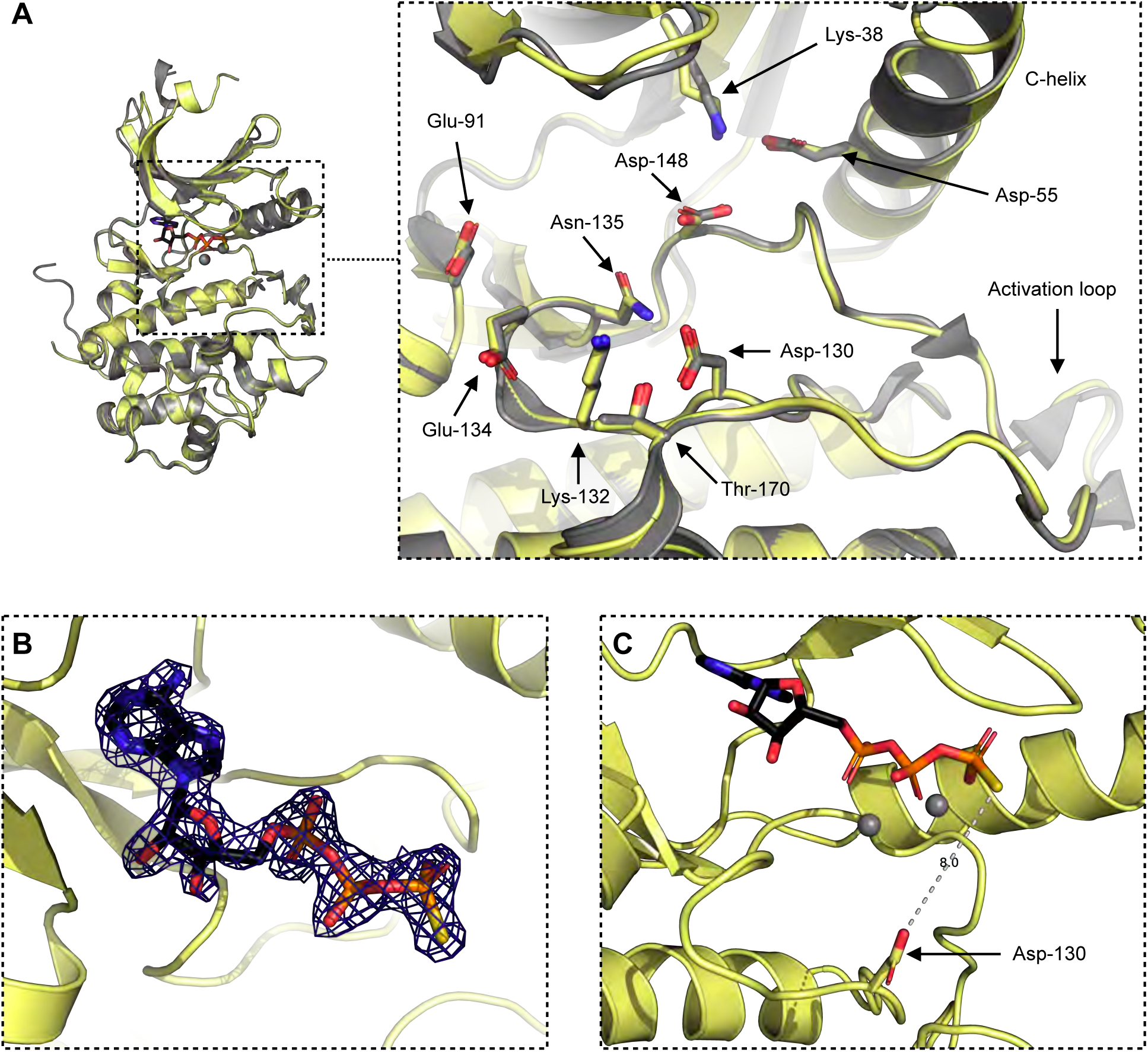
**a)** Superposition of the ATPyS bound CHK1 structure (yellow) with an apo CHK1 structure 1IA8 (grey), both the overall arrangement of the N and C kinase lobes, and the details of key active site residues are identical between the two models. **b)** 2mFo-DFc map showing electron density for ATPyS in the structure. **c)** Details of the distance between the catalytic Asp-130 residue and the γ-phosphate of the ATPγS.

### Crystal structure of CHK1 bound to CLASPIN

A further trial where the kinase domain was mixed with the CLASPIN peptide and the broad-spectrum ATP-competitive kinase inhibitor staurosporine, produced a structure where both the inhibitor and the peptide were present. There are two independent copies of the kinase domain in the asymmetric unit of this crystal lattice, and while both of the CHK1 molecules are able accommodate the CLASPIN peptide, in one of the chains the peptide is constrained by the lattice and adopts an artefactual conformation where the peptide interacts with multiple CHK1 molecules. The CLASPIN peptides bound to the other chain adopts a position and conformation situated between, and making interactions with, the C-helix and the activation loop of a single kinase domain (**Figure 4a**) with density for the peptide visible in an omit map (**Figure 4b**).

**FIGURE 4.**
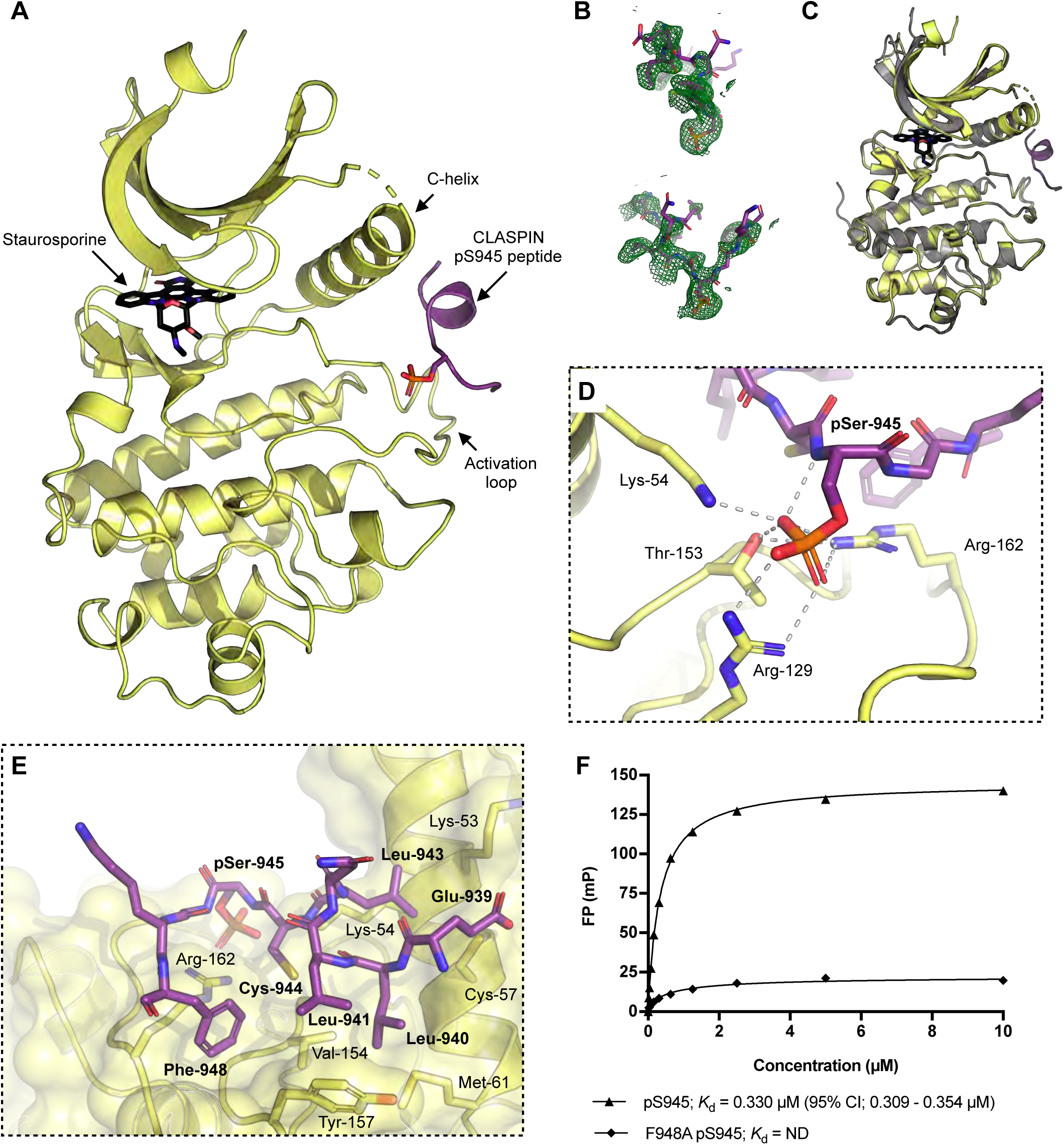
**a)** Structure of a CLASPIN pS945 peptide (purple) bound to CHK1-KD (yellow) with staurosporine bound in the active site. **b)** A 2mFo-DFc omit map showing density for the CLASPIN peptide in the structure **c)** Superposition of the CLASPIN pS945 CHK1 structure (yellow) with a Staurosporine bound CHK1 structure 1NVR (grey). The overall arrangement of the N and C kinase lobes are identical between the two models. **d)** Details for the interaction with the phosphate group of the bound CLASPIN peptide, residues in the peptide are labelled in bold. **e)** Details of the hydrophobic interactions made between the kinase domain and the CLASPIN peptide. **f)** Fluorescence polarisation experiments show binding between the CLASPIN peptide and CHK1 can be abrogated by a mutation in the CLASPIN sequence.

Notably the overall positioning of the kinase domain lobes to each other, and the active site architecture, is the same as previously determined apo or inhibitor bound structures (**Figure 4c**), with an overall RMSD of 0.252Å for all protein atoms from a staurosporine-bound structure (Zhao et al., 2002). This suggests that CLASPIN binding does not play a significant role in aligning the active site residues into a catalytically active conformation.

The phosphorylated state of the CLASPIN peptide is recognised by the side chains of CHK1 Lys-54, Arg-129, Thr-153 and Arg-162 co-ordinating the phosphate group of CLASPIN pSer-945, with additional H-bond interactions between the mainchain carbonyl of CLASPIN pSer-945 and the sidechain of CHK1 Arg-162 (**Figure 4d**). Specificity for the CLASPIN sequence beyond the phosphorylation, is provided by a series of predominantly hydrophobic interactions (**Figure 4e**). Leu-943 of CLASPIN sits between the side chains of CHK1 Lys-53, Lys-54 and Cys 57 while CLASPIN Leu-940, Leu-941 and Phe-948 protrude into a hydrophobic groove on the surface of the kinase domain lined by the side chains of CHK1 Met-61, Val-154, Tyr-157 and Arg-162.

In order to validate the CLASPIN binding mode seen in the crystal structure, a peptide with a point mutation in the phenylalanine conserved between the three motifs, and shown from the crystal structure to be important for the interaction, was synthesised and its binding to CHK1-KD measured using fluorescence polarisation. The mutation was found to significantly lower the affinity of the interaction in the FP assay (**Figure 4f**).

### CLASPIN binding is uncoupled from CHK1 regulation

The position of the CLASPIN binding site on CHK1 that our study reveals, is distant from the proposed CHK1-KA1 domain binding site based on models derived from the MARK1 kinase structure (Emptage et al., 2018) (**Figure 5a**), but in a similar location to the cyclin binding site in CDKs (Russo et al., 1996). This suggests that CLASPIN binding would be unaffected by the KA1 domain-regulated activation state of CHK1. To test this, we purified full-length CHK1, treated it with λ-phosphatase to remove any activating phosphorylation, and measured its affinity for the CLASPIN-phosphopeptide in a fluorescence polarisation assay (**Figure 5b**). As predicted, the CLASPIN peptide still bound KA1-autoinhibited CHK1, but with two-fold lower affinity than for CHK1-KD, indicating some negative cooperativity between KA1-domain and CLASPIN binding. However, the relative weakness of the effect suggests that it is unlikely to be biologically significant.

**FIGURE 5.**
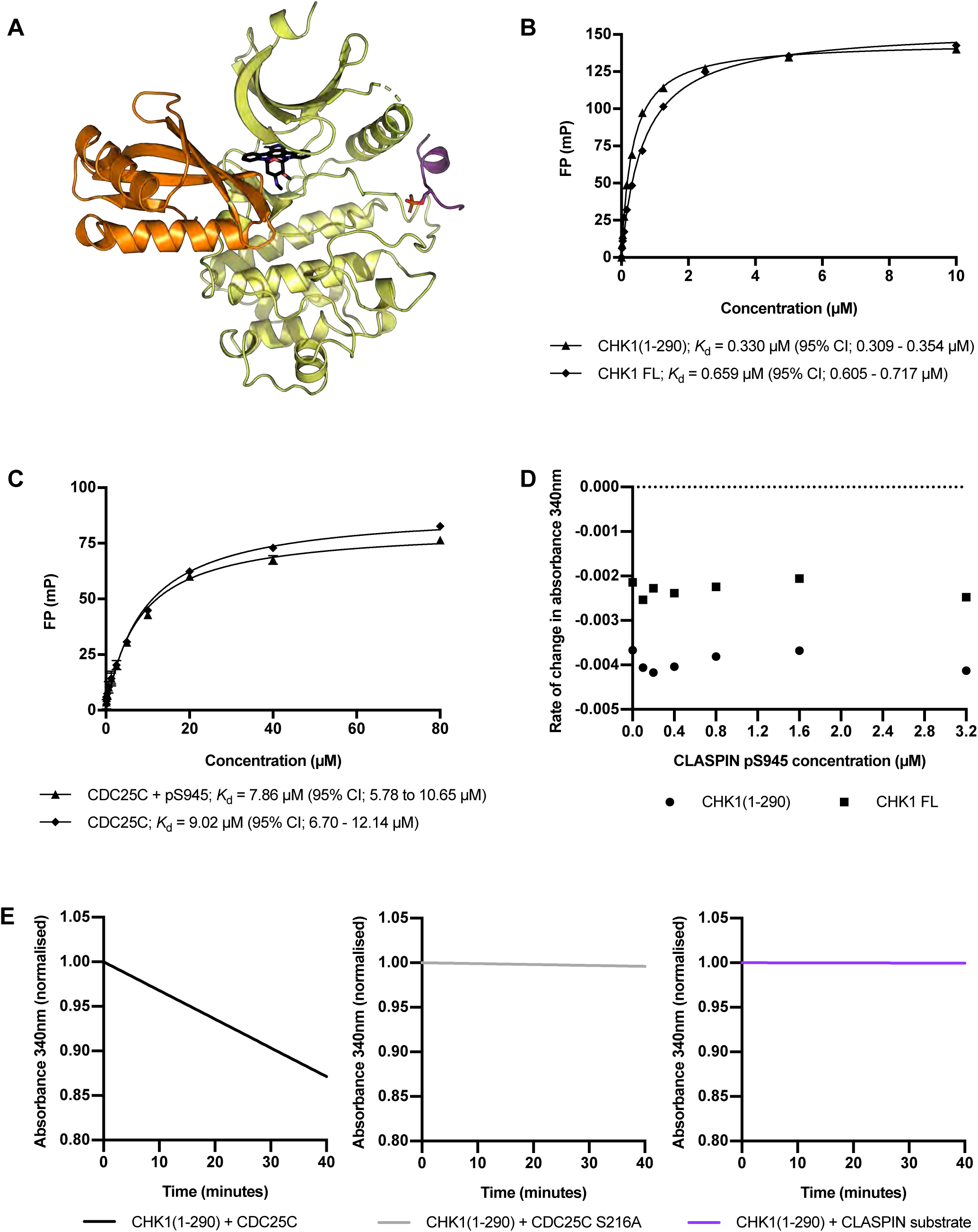
**a)** Model for the interaction of the KA1 domain (orange) with the kinase domain (yellow) for CHK1 produced by superposing both the KA1 domain, and the CLASPIN bound kinase domain, on the KA1-autoinhibited MARK1 kinase structure 6C9D.Fluorescence polarisation experiments showing CLASPIN peptide interactions with full length CHK1 or a CHK1 construct consisting of only the kinase domain. The CLASPIN peptide binds with comparable affinity to both constructs. **b)** Fluorescence polarisation experiments showing CDC25C substrate peptide interactions with a CHK1 construct consisting of only the kinase domain, in the presence or absence of the CLASPIN pS945 peptide. The presence of the CLASPIN peptide has no significant effect on the affinity of the kinase for the substrate peptide. **c)** NADH-coupled assay showing ATP turnover by either full length CHK1 or a CHK1 construct consisting of only the kinase domain in the presence of increasing concentrations of the CLASPIN pS945 peptide. Taken together with **c)** these data show that CLASPIN binding has no allosteric effect on CHK1 kinase function. **d)** NADH-coupled ATPase assay for CHK1-KD incubated with a CDC25C substrate, mutant S216A CDC25C substrate and CLASPIN S945 substrate. These data show that the CLASPIN peptide itself is not a substrate for CHK1 kinase, contrary to previous suggestions (Chini and Chen, 2006).

### CLASPIN binding does not affect CHK1 activity

The coincidence of the CLASPIN-binding site in CHK1 with cyclin-binding to CDKs suggests that CLASPIN, like cyclins, might modulate the affinity of the kinase for its substrate and/or affect its catalytic activity. To test this, we used fluorescence polarisation to determine the affinity of a CDC25C substrate peptide for CHK1-KD in the presence or absence of the pS945 CLASPIN peptide, but saw no significant difference (**Figure 5c**), suggesting that the CHK1-CLASPIN interaction does not modulate the affinity of CHK1-KD for substrate.

To determine whether CLASPIN binding has an allosteric effect on CHK1 kinase activity, we used an NADH-coupled ATPase assay to determine the rate of ATP turnover in the presence of a substrate peptide corresponding to the CHK1 phosphorylation site in CDC25C. Under the conditions tested the rate of the reaction was 2.25 mol min^−1^ mol^−1^ for the full-length kinase and 7.73 mol min^−1^ mol^−1^ for the kinase domain alone. For both full-length CHK1 and CHK1-KD, the addition of the CLASPIN pS945 peptide had no effect on the rate of ATP turnover of the reaction in the presence of the CDC25C peptide (**Figures 5d**). In order to show that the ATP turnover measured was the result of the phosphorylation of Ser-216 in the CDC25C peptide, rather than some other activity, a CLASPIN substrate peptide in which Ser-216 was replaced with an alanine was used, which showed no activity in the assay. We also looked at whether the CLASPIN peptide itself could be phosphorylated by CHK1 as has been suggested, (Chini and Chen, 2006), but saw no stimulation of ATP turnover by CHK1 in the presence of the unphosphorylated S945 CLASPIN peptide (**Figures 5e**).

## DISCUSSION

The results presented here provide a reliable model for the nucleotide bound state of CHK1-KD and reveal the molecular basis for the phosphorylation dependent interaction between CHK1 and CLASPIN. Our data show that CLASPIN binding has no significant direct effect on the regulation or kinase activity of CHK1, but suggest that CLASPIN acts as a pure scaffold, facilitating the recruitment of CHK1 into a larger complex in which it can be activated through phosphorylation by ATR, downstream of DNA damage.

The interactions of the bound CLASPIN motif with CHK1 explains the pattern of small hydrophobic residues at positions −1, −2, −4 and −5 relative to the phosphorylated Ser/Thr, and a glycine at +1 and phenylalanine at +3, all of which are very strongly conserved in the three repeats of this motif that occur throughput the vertebrates. As our structural and biochemical data show that a single copy of this motif is sufficient for a high-affinity interaction with a unique site on CHK1, the apparent biological need for a triple tandem array of these motifs in CLASPIN is far from clear. Pull-down studies of *Xenopus* CLASPIN, in which the third repeat is less strongly conserved, indicated some possible cooperativity in CHK1 binding between the first two sites (Kumagai and Dunphy, 2003), favouring binding of multiple CHK1 molecules to a single CLASPIN. Focal accumulation of CHK1 at sites of ATR activation can be observed in cells following DNA damage (Bigot et al., 2019), and recruitment of multiple CHK1 molecules by each CLASPIN might be a means of generating a high concentration of CHK1 activity at sites of DNA damage.

Clustering of CHK1 molecules by CLASPIN could also play a role in the second stage of the CHK1 kinase activation process, which requires autophosphorylation at two consensus CHK1-phosphorylation motifs and at least one non-consensus site within the KA1 domain, subsequent to phosphorylation by ATR (Gong et al., 2018; Kasahara et al., 2010; Okita et al., 2012). Autophosphorylation mediated by phosphorylation-dependent clustering downstream of phosphorylation by ATM also underlies activation of the other checkpoint kinase CHK2 (Ahn et al., 2002; Oliver et al., 2006). However, the two processes are mechanistically distinct, as CHK1 autophosphorylation occurs in an ancillary domain and may be partly intramolecular (Okita et al., 2012), while *trans*-autophosphorylation of CHK2 and related kinases occurs within the kinase domain itself and is strictly intermolecular (Oliver et al., 2007; Pike et al., 2008).

An alternative function for the tandem motifs of CLASPIN is as an internal scaffold, whereby prior phosphorylation of one motif that engages with the CLASPIN-binding site on CHK1, facilitates binding of an adjacent unphosphorylated motif to the active site of CHK1 as a substrate, by increasing its local concentration. Phosphorylation of CLASPIN by CHK1 *in vitro* has been reported (Chini and Chen, 2006), but subsequent studies show that CHK1 is not required for CLASPIN phosphorylation *in vivo* (Bennett et al., 2008), where this function is fulfilled by CK1γ1 (Meng et al., 2011) and CDC7 (Yang et al., 2019). Our *in vitro* data show that an isolated unphosphorylated CLASPIN motif displays negligible affinity for CHK1 and no significant activity as a kinase substrate. While we cannot completely rule out the possibility that CHK1 might phosphorylate a longer CLASPIN peptide spanning two or more motifs, where one is already phosphorylated, the lack of even a weak affinity or activity for the isolated unphosphorylated CLASPIN CHK1-binding motif argues against this.

### CLASPIN utilises a conserved phosphate-binding pocket

The site occupied by the phosphorylated Ser/Thr in CLASPINs CHK1-binding motifs is a phosphate binding pocket that serves at least four different functions across the different kinases within which it is evolutionarily conserved (**Figure 6**). In all cases the residues that coordinate a bound phosphate group are highly conserved, but the rest of the binding site that provides the larger recognition process in each case, is not. The most widely observed role for this site, is to bind to the side-chain of a phosphorylated threonine within the activation segment of the kinase, and bridge it to the C-helix within the N-terminal lobe (Endicott et al., 2012). Thus, for example, in CDK2, the side chains of Arg-50, Arg-126 and Arg-150 co-ordinate the phosphate group of pThr-160 of CDK2 (**Figure 6b**), thereby stabilising the active conformation of the enzyme (Bao et al., 2011). The equivalent site in GSK3β (Arg-96, Arg-180 and Lys-205) functions in the specific recognition of ‘primed’ substrates by coordinating the phosphate group on already phosphorylated residues within the substrate, positioned four residues downstream of the residue being phosphorylated, and thereby greatly enhances their affinity for the kinase and the efficiency of their phosphorylation (Dajani et al., 2001). This site is also used to regulate GSK3β kinase activity by facilitating binding of phosphorylated residues in inhibitory sequences, such as pSer-9 in the autoinhibitory N-terminus of GSK3β itself, or pSer-1607 in low-density lipoprotein receptor-like protein 6 (LRP6) (Stamos et al., 2014) (**Figure 6c**), so that the upstream sequence occupies the active site and prevents substrate access. The structure of the CLASPIN-CHK1 complex described here adds a hitherto unknown role for this site, in providing a phosphorylation-dependent scaffold interaction that facilitates recruitment of the kinase to a specific cellular location.

**FIGURE 6.**
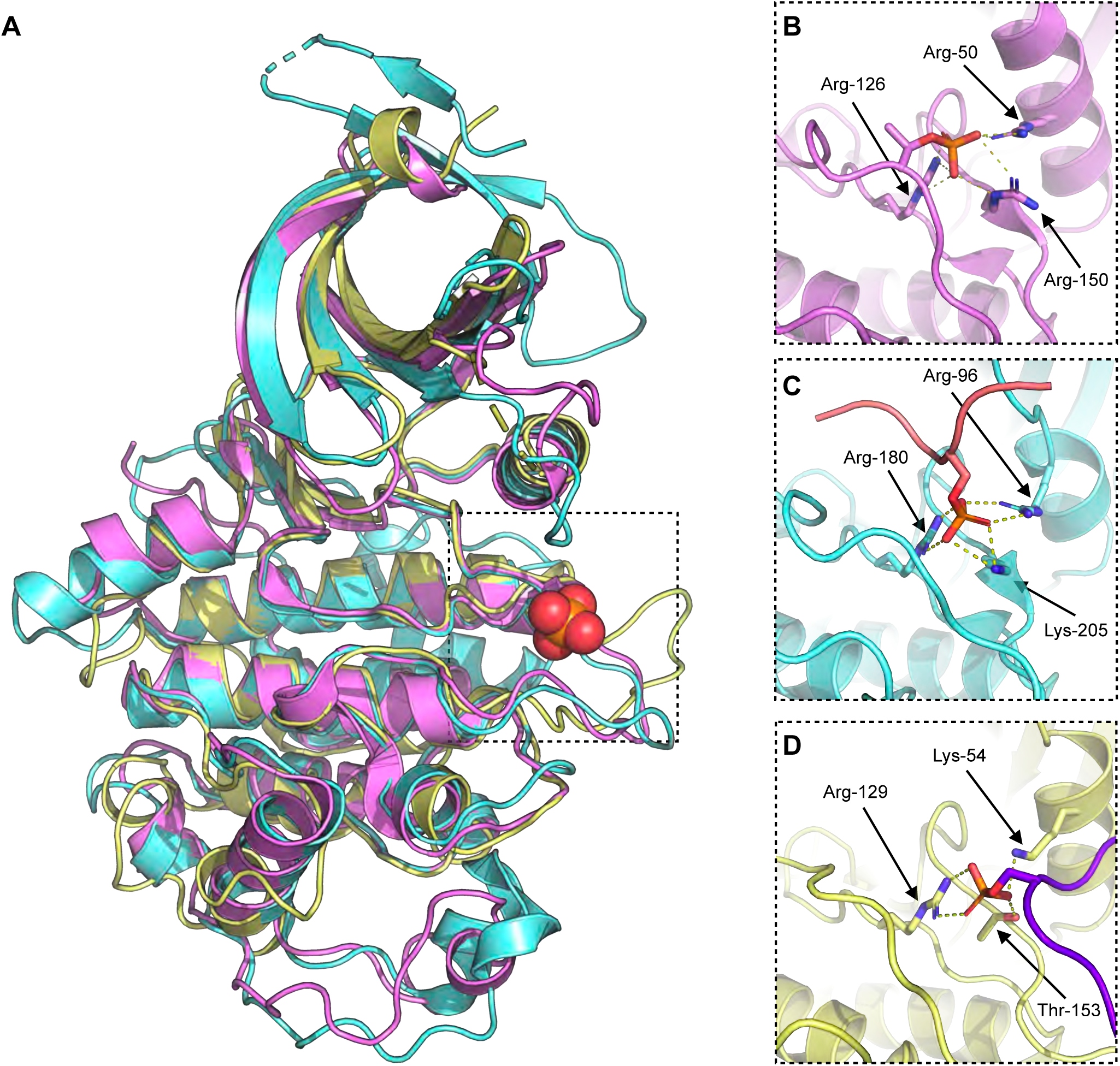
**a)** Superposition of the kinase domains and interacting regions of CDK2 (pink), GSK3β (cyan), CHK1 (yellow) showing location of conserved phosphate binding pocket, marked with a phosphate group from the CHK1-CLASPIN structure presented here. **b-d)** Details of the interaction with phosphorylated residues in CDK2, GSK3β and CHK1 respectively. The LRP6 inhibitor peptide bound to GSK3β is shown in pink and the CLASPIN scaffold peptide bound to CHK1 in purple.

**Table 1.** Data collection and refinement statistics. Statistics for the highest-resolution shell are shown in parentheses.

|  | CHK1 ATPyS | CHK1 Staurosporine CLASPIN |
| --- | --- | --- |
| Wavelength | 0.96864 | 0.96864 |
| Resolution range | 42.6 - 1.93 (1.999 - 1.93) | 96.49 - 1.81 (1.842 - 1.81) |
| Space group | P 21 21 21 | P 1 21 1 |
| Unit cell | 65.62 89.27 112.03 90 90 90 | 60.34 64.45 97.08 90 96.32 90 |
| Total reflections | 195490 (15117) | 191511 (5685) |
| Unique reflections | 49422 (4447) | 65068 (2602) |
| Multiplicity | 4.0 (3.4) | 2.9 (2.2) |
| Completeness (%) | 98.35 (89.84) | 96.2 (76.5) |
| Mean I/sigma(I) | 5.57 (0.79) | 8.8 (0.4) |
| Wilson B-factor | 28.60 | 29.03 |
| R-merge | 0.1555 (1.313) | 0.063 (1.855) |
| R-meas | 0.1795 (1.54) | 0.076 (2.365) |
| R-pim | 0.08786 (0.7813) | 0.043 (1.445) |
| CC1/2 | 0.992 (0.336) | 0.998 (0.161) |
| Reflections used in refinement | 49384 (4447) | 51946 (377) |
| Reflections used for R-free | 2410 (224) | 2513 (22) |
| R-work | 0.1967 (0.3214) | 0.1901 (0.3345) |
| R-free | 0.2417 (0.3609) | 0.2353 (0.4285) |
| CC(work) | 0.951 (0.648) | 0.932 (0.643) |
| CC(free) | 0.926 (0.580) | 0.933 (0.752) |
| Number of non-hydrogen atoms | 4903 | 4982 |
| macromolecules | 4385 | 4569 |
| ligands | 83 | 99 |
| solvent | 435 | 314 |
| Protein residues | 538 | 561 |
| RMS(bonds) | 0.005 | 0.004 |
| RMS(angles) | 0.78 | 0.73 |
| Ramachandran favored (%) | 97.19 | 96.37 |
| Ramachandran allowed (%) | 2.81 | 3.09 |
| Ramachandran outliers (%) | 0.00 | 0.54 |
| Rotamer outliers (%) | 0.00 | 1.21 |
| Clashscore | 3.59 | 5.25 |
| Average B-factor | 36.48 | 40.63 |
| macromolecules | 35.97 | 40.67 |
| ligands | 41.54 | 32.25 |
| solvent | 40.66 | 42.75 |

## EXPERIMENTAL PROCEDURE

### Protein Expression and Purification

Recombinant full length CHK1, or kinase domain alone (CHK1-KD), was produced by infecting Sf9 cells with a baculovirus coding for a construct containing STREP-(3C)-CHK1-HIS or STREP-(3C)-CHK1(1-290)-HIS respectively. Cell pellets were re-suspended in lysis buffer containing 50mM HEPES pH 7.5, 200mM NaCl, 0.5mM TCEP, and supplemented with 10 U DNASE Turbo, then disrupted by sonication, and the resulting lysate clarified by centrifugation at 40,000 ⨯ g for 60 minutes at 4°C. The supernatant was applied to a 5ml HiTrap TALON crude column (GE Healthcare, Little Chalfont, UK), washed first with buffer containing 50mM HEPES pH 7.5, 500mM NaCl, 0.5mM TCEP, followed lysis buffer supplemented with 10 mM 10mM imidazole, with any retained protein then eluted by application of the same buffer but now supplemented with 250mM imidazole. The eluted protein was diluted with lysis buffer before application to a 5ml HiTrap STREP column (GE Healthcare, Little Chalfont, UK), washed with lysis buffer and eluted using buffer supplemented with 2 mM Desthiobiotin.

For FP experiments, a Superdex 75 16/60 size exclusion column (GE Healthcare) was used to purify the protein to homogeneity in 25mM HEPES pH 7.5, 200mM NaCl, 1mM EDTA, 0.25mM TCEP, 0.02% (v/v) Tween-20.

For crystallographic studies, the N-terminal STREP tag was removed by incubation with GST-3C protease (in house) for 12 hours at 4°C. A Superdex 75 16/60 size exclusion column (GE Healthcare) was used to purify the kinase domain to homogeneity in 10mM HEPES pH 7.5, 200mM NaCl, 0.5mM TCEP.

### Fluorescence Polarisation Experiments

Fluorescein-labelled (Flu-pT916 Flu-GGMDELLDLC(pT)GKFTSQ, Flu-pS945 Flu-GGMEELLNLC(pS)GKFTSQ and Flu-pS982 Flu-GGMEEALALC(pS)GSFPTD) or CY5-labelled (Cy5-CDC25C-S216 Cy5-CGYGGLYRSPSMPENLNRPRLK) peptides (Peptide Protein Research Ltd, Bishops Waltham, UK) at a concentration of 100nM, or for the double peptide experiment 200 nM Cy5-CDC25C and 1000 nM CLASPIN pS945, were incubated at room temperature with increasing concentrations of CHK1-KD, or lambda phosphatase treated full length CHK1 in 25mM HEPES pH 7.5, 200mM NaCl, 1mM EDTA, 0.25mM TCEP, 0.02% (v/v) Tween-20 in a black 96-well polypropylene plate (VWR, Lutterworth, UK). Incubation with Lambda phosphatase (NEB, Hitchin, UK) supplemented with 1 mM MnCl2 was used to remove the phosphorylation on the peptides and Staurosporin (NEB, Hitchin, UK) to occupy the active site pocket. Fluorescence polarisation was measured in a POLARstar Omega multimode microplate reader (BMG Labtech GmbH, Offenburg, Germany). Binding curves represent the mean of 3 independent experiments, with error bars of 1 standard deviation. All data were fitted by non-linear regression, to a one site – specific binding model in Prism 6 for Mac OS X (v 6.0d, GraphPad Software) in order to calculate the reported disassociation constants (*K*_d_).

### Crystallography

The pure CHK1-KD was mixed with a fifty molar excess of ATPyS (Sigma-Aldrich, Gillingham, United Kingdom) and a five molar excess of both a substrate peptide corresponding to residues 210-227 of CDC25 (GLYRSPSMPENLNRPRLK) and a pS945 CLASPIN peptide (MEELLNLC(pS)GKFT(pS)QD) (Peptide Protein Research Ltd, Bishops Waltham, UK) prior to concentration to 8 mg/ml for use in crystallisation trials. Crystals grew in condition PACT H11 (200 mM Sodium citrate tribasic dihydrate, 100 mM Bis-Tris propane pH 8.5 and 20% w/v PEG 3350) and following further optimisation of this condition single crystals were looped, soaked and cryoprotected in well solution supplemented with 30 % Ethylene Glycol before flash freezing in liquid nitrogen.

Staurosporin (NEB, Hitchin, UK) and the pS945 CLASPIN peptide (MEELLNLC(pS)GKFT(pS)QD) (Peptide Protein Research Ltd, Bishops Waltham, UK) were mixed with the pure CHK1-KD at two and five molar excess respectively prior to concentration to 8 mg/ml for use in crystallisation trials. Crystals grew in condition PACT F11 (200 mM Sodium citrate tribasic dihydrate, 100 mM Bis-Tris propane pH 6.5 and 20% w/v PEG 3350) and following further optimisation of this condition single crystals were looped, soaked and cryoprotected in well solution supplemented with 30 % Ethylene Glycol and 10 mM peptide before flash freezing in liquid nitrogen. Data were collected on beamline I24 at the Diamond Synchrotron Lightsource and the structure was determined using PHASER (McCoy, 2017) to perform molecular replacement with PDB 1IA8 as a search model before refinement using the PHENIX software package (Headd et al., 2012).

### ATPase assay

CHK1, to a concentration of 100 nM for the kinase domain or 200 nM for full length CHK1, was added to reaction mixtures containing Pyruvate Kinase and Lactic Dehydrogenase enzymes (Sigma-Aldrich, Gillingham, United Kingdom) with 0.010-0.016, and 0.015-0.023 units of activity respectively in 25 mM HEPES pH 7.5, 150 mM NaCl, 0.5mM TCEP, 0.5 mM PEP, 0.5 mM NADH, 1 mM ATP with 0.5 mM CDC25C (210-227) peptide (GLYRSPSMPENLNRPRLK) or CDC25C S216A (210-227) peptide (GLYRSPSMPENLNRPRLK) and with or without CLASPIN pS945 peptide (MEELLNLC(pS)GKFTSQ) (Peptide Protein Research Ltd, Bishops Waltham, UK) at different concentrations in a Thermoscientific 96 well UV microplate (Fisher scientific, Loughborough, UK). The plates were incubated at 25 °C and absorbance at 340 nm was measured in a POLARstar Omega multimode microplate reader (BMG Labtech GmbH, Offenburg, Germany) at intervals of 30 seconds.

## Acknowledgements

We thank Mark Roe for assistance with X-ray data collection and both Lihong Zhou and Raquel Arribas for help with baculovirus protein expression. We thank Dr Neil Kad (Uni. of Kent) for details of the NADH-coupled ATPase assay. We are grateful to the Diamond Light Source Ltd., Didcot, UK, for access to synchrotron radiation and to the Wellcome Trust for support for X-ray diffraction facilities at the University of Sussex. This work was supported by Cancer Research UK Programme Grants C302/A14532 and C302/A24386 (A.W.O and L.H.P.).

## Author Contributions

Conceptualization : M.D., A.W.O., L.H.P.; Methodology : M.D., A.W.O., L.H.P.; Validation : M.D., A.W.O., L.H.P.; Formal Analysis : M.D., A.W.O., L.H.P.; Investigation : All Authors; Writing – Original Draft : M.D.; Writing – Review & Editing :M.D., A.W.O., L.H.P.; Visualisation M.D.; Supervision : L.H.P., A.W.O.; Funding Acquisition : L.H.P., A.W.O.

## Data availability

Structure Factors and refined atomic coordinates have been deposited in the PDB with accession codes : 7AKM and 7AKO.

---

**Supplementary Figure 5.**
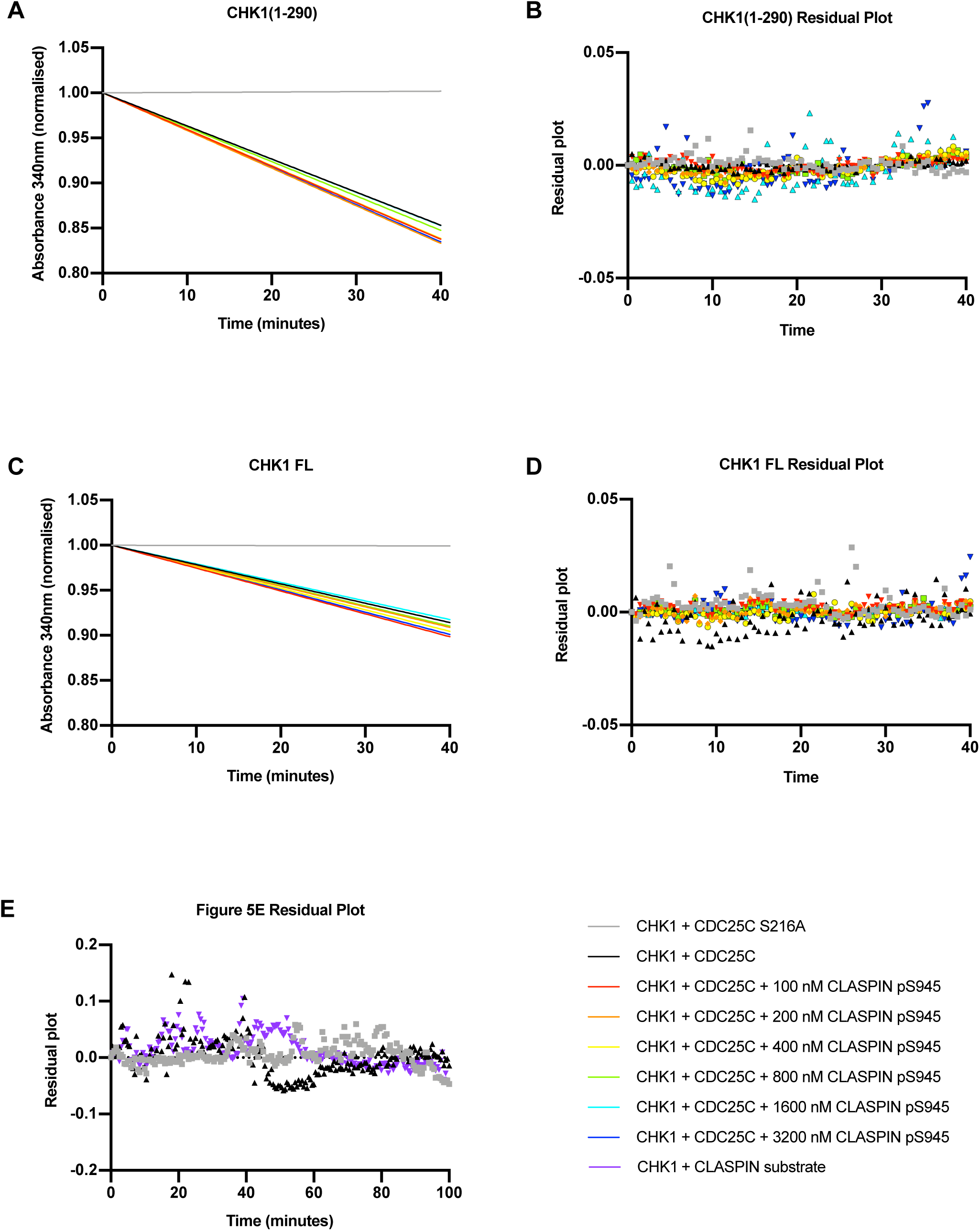

## REFERENCES

Ahn, J.Y., Li, X., Davis, H.L., and Canman, C.E. (2002). Phosphorylation of threonine 68 promotes oligomerization and autophosphorylation of the Chk2 protein kinase via the forkhead-associated domain. J Biol Chem 277, 19389–19395.

Bao, Z.Q., Jacobsen, D.M., and Young, M.A. (2011). Briefly bound to activate: transient binding of a second catalytic magnesium activates the structure and dynamics of CDK2 kinase for catalysis. Structure 19, 675–690.

Bennett, L.N., Larkin, C., Gillespie, D.A., and Clarke, P.R. (2008). Claspin is phosphorylated in the Chk1-binding domain by a kinase distinct from Chk1. Biochem Biophys Res Commun 369, 973–976.

Bigot, N., Day, M., Baldock, R.A., Watts, F.Z., Oliver, A.W., and Pearl, L.H. (2019). Phosphorylation-mediated interactions with TOPBP1 couple 53BP1 and 9-1-1 to control the G1 DNA damage checkpoint. Elife 8.

Chen, P., Luo, C., Deng, Y., Ryan, K., Register, J., Margosiak, S., Tempczyk-Russell, A., Nguyen, B., Myers, P., Lundgren, K., et al. (2000). The 1.7 A crystal structure of human cell cycle checkpoint kinase Chk1: implications for Chk1 regulation. Cell 100, 681–692.

Chini, C.C., and Chen, J. (2006). Repeated phosphopeptide motifs in human Claspin are phosphorylated by Chk1 and mediate Claspin function. J Biol Chem 281, 33276–33282.

Clarke, C.A., and Clarke, P.R. (2005). DNA-dependent phosphorylation of Chk1 and Claspin in a human cell-free system. Biochem J 388, 705–712.

Dajani, R., Fraser, E., Roe, S.M., Young, N., Good, V., Dale, T.C., and Pearl, L.H. (2001). Crystal structure of glycogen synthase kinase 3 beta: structural basis for phosphate-primed substrate specificity and autoinhibition. Cell 105, 721–732.

Emptage, R.P., Lemmon, M.A., Ferguson, K.M., and Marmorstein, R. (2018). Structural Basis for MARK1 Kinase Autoinhibition by Its KA1 Domain. Structure 26, 1137–1143 e1133.

Emptage, R.P., Schoenberger, M.J., Ferguson, K.M., and Marmorstein, R. (2017). Intramolecular autoinhibition of checkpoint kinase 1 is mediated by conserved basic motifs of the C-terminal kinase-associated 1 domain. J Biol Chem 292, 19024–19033.

Endicott, J.A., Noble, M.E., and Johnson, L.N. (2012). The structural basis for control of eukaryotic protein kinases. Annu Rev Biochem 81, 587–613.

Gong, E.Y., Hernandez, B., Nielsen, J.H., Smits, V.A.J., Freire, R., and Gillespie, D.A. (2018). Chk1 KA1 domain auto-phosphorylation stimulates biological activity and is linked to rapid proteasomal degradation. Sci Rep 8, 17536.

Gong, E.Y., Smits, V.A.J., Fumagallo, F., Piscitello, D., Morrice, N., Freire, R., and Gillespie, D.A. (2015). KA1-targeted regulatory domain mutations activate Chk1 in the absence of DNA damage. Sci Rep 5, 10856.

Headd, J.J., Echols, N., Afonine, P.V., Grosse-Kunstleve, R.W., Chen, V.B., Moriarty, N.W., Richardson, D.C., Richardson, J.S., and Adams, P.D. (2012). Use of knowledge-based restraints in phenix.refine to improve macromolecular refinement at low resolution. Acta Crystallogr D Biol Crystallogr 68, 381–390.

Iyer, D.R., and Rhind, N. (2017). The Intra-S Checkpoint Responses to DNA Damage. Genes (Basel) 8.

Jeong, S.Y., Kumagai, A., Lee, J., and Dunphy, W.G. (2003). Phosphorylated claspin interacts with a phosphate-binding site in the kinase domain of Chk1 during ATR-mediated activation. J Biol Chem 278, 46782–46788.

Kasahara, K., Goto, H., Enomoto, M., Tomono, Y., Kiyono, T., and Inagaki, M. (2010). 14-3-3gamma mediates Cdc25A proteolysis to block premature mitotic entry after DNA damage. EMBO J 29, 2802–2812.

Kumagai, A., and Dunphy, W.G. (2000). Claspin, a novel protein required for the activation of Chk1 during a DNA replication checkpoint response in Xenopus egg extracts. Mol Cell 6, 839–849.

Kumagai, A., and Dunphy, W.G. (2003). Repeated phosphopeptide motifs in Claspin mediate the regulated binding of Chk1. Nat Cell Biol 5, 161–165.

Kumagai, A., Kim, S.M., and Dunphy, W.G. (2004). Claspin and the activated form of ATR-ATRIP collaborate in the activation of Chk1. J Biol Chem 279, 49599–49608.

Lindsey-Boltz, L.A., Sercin, O., Choi, J.H., and Sancar, A. (2009). Reconstitution of human claspin-mediated phosphorylation of Chk1 by the ATR (ataxia telangiectasia-mutated and rad3-related) checkpoint kinase. J Biol Chem 284, 33107–33114.

McCoy, A.J. (2017). Acknowledging Errors: Advanced Molecular Replacement with Phaser. Methods Mol Biol 1607, 421–453.

Meng, Z., Capalbo, L., Glover, D.M., and Dunphy, W.G. (2011). Role for casein kinase 1 in the phosphorylation of Claspin on critical residues necessary for the activation of Chk1. Mol Biol Cell 22, 2834–2847.

Niida, H., Katsuno, Y., Banerjee, B., Hande, M.P., and Nakanishi, M. (2007). Specific role of Chk1 phosphorylations in cell survival and checkpoint activation. Mol Cell Biol 27, 2572–2581.

Okita, N., Minato, S., Ohmi, E., Tanuma, S., and Higami, Y. (2012). DNA damage-induced CHK1 autophosphorylation at Ser296 is regulated by an intramolecular mechanism. FEBS Lett 586, 3974–3979.

Oliver, A.W., Knapp, S., and Pearl, L.H. (2007). Activation segment exchange: a common mechanism of kinase autophosphorylation? Trends Biochem Sci 32, 351–356.

Oliver, A.W., Paul, A., Boxall, K.J., Barrie, S.E., Aherne, G.W., Garrett, M.D., Mittnacht, S., and Pearl, L.H. (2006). Trans-activation of the DNA-damage signalling protein kinase Chk2 by T-loop exchange. EMBO J 25, 3179–3190.

Pike, A.C., Rellos, P., Niesen, F.H., Turnbull, A., Oliver, A.W., Parker, S.A., Turk, B.E., Pearl, L.H., and Knapp, S. (2008). Activation segment dimerization: a mechanism for kinase autophosphorylation of non-consensus sites. EMBO J 27, 704–714.

Russo, A.A., Jeffrey, P.D., and Pavletich, N.P. (1996). Structural basis of cyclin-dependent kinase activation by phosphorylation. Nat Struct Biol 3, 696–700.

Smits, V.A., and Gillespie, D.A. (2015). DNA damage control: regulation and functions of checkpoint kinase 1. FEBS J 282, 3681–3692.

Stamos, J.L., Chu, M.L., Enos, M.D., Shah, N., and Weis, W.I. (2014). Structural basis of GSK-3 inhibition by N-terminal phosphorylation and by the Wnt receptor LRP6. Elife 3, e01998.

Wright, P.E., and Dyson, H.J. (2015). Intrinsically disordered proteins in cellular signalling and regulation. Nat Rev Mol Cell Biol 16, 18–29.

Yang, C.C., Kato, H., Shindo, M., and Masai, H. (2019). Cdc7 activates replication checkpoint by phosphorylating the Chk1-binding domain of Claspin in human cells. Elife 8.

Zhao, B., Bower, M.J., McDevitt, P.J., Zhao, H., Davis, S.T., Johanson, K.O., Green, S.M., Concha, N.O., and Zhou, B.B. (2002). Structural basis for Chk1 inhibition by UCN-01. J Biol Chem 277, 46609–46615.

